# Molecular Graph Enhanced Transformer for Retrosynthesis Prediction

**DOI:** 10.1101/2020.03.05.979773

**Authors:** Kelong Mao, Peilin Zhao, Tingyang Xu, Yu Rong, Xi Xiao, Junzhou Huang

**Affiliations:** Tsinghua University; Tencent AI Lab

**Author notes:** }. This work is done when Kelong Mao works as an intern in Tencent AI Lab.

## Abstract

With massive possible synthetic routes in chemistry, retrosynthesis prediction is still a challenge for researchers. Recently, retrosynthesis prediction is formulated as a Machine Translation (MT) task. Namely, since each molecule can be represented as a Simplified Molecular-Input Line-Entry System (SMILES) string, the process of retrosynthesis is analogized to a process of language translation from the product to reactants. However, the MT models that applied on SMILES data usually ignore the information of natural atomic connections and the topology of molecules. To make more chemically plausible constrains on the atom representation learning for better performance, in this paper, we propose a Graph Enhanced Transformer (GET) framework, which adopts both the sequential and graphical information of molecules. Four different GET designs are proposed, which fuse the SMILES representations with atom embeddings learned from our improved Graph Neural Network (GNN). Empirical results show that our model significantly outperforms the vanilla Transformer model in test accuracy.

## 1 Introduction

Retrosynthesis prediction aims to predict a set of suitable reactants that can synthesize the desired molecule via a series of reactions. It pushes forward an immense influence in agriculture, medical treatment, drug discovery and so on. However, the retrosynthesis prediction is challenging since there are massive possible synthetic routes available and it is often difficult to navigate the direction of retrosynthesis process. Indeed, each bond in the target molecule may represent a possible retrosynthetic disconnection, leading to a vast space of possible starting materials. Besides, the difference between two synthetic routes may be subtle, which usually depends on the global molecular structures. Actually, planning a proper retrosynthetic route for a complex molecule is also a tough work even for the professional chemists.

One of the prevailing methods is to deem the retrosynthesis prediction as a machine translation task. This analogy is comprehensible since every molecule has a unique text representation named SMILES [Weininger, 1988]. In this case, given a target molecule written in SMILES notation, the retrosynthesis prediction is just to predict a string of SMILES which represents the reactants. Based on this idea, [Liu *et al.*, 2017] first applied LSTM with attention mechanism in retrosynthesis prediction and achieved comparable performance. Whereafter, many works [Karpov *et al.*, 2019; Zheng *et al.*, 2019; Lin *et al.*, 2019; Lee *et al.*, 2019] tried to employ Transformer [Vaswani *et al.*, 2017], a more powerful Sequence-to-Sequence(Seq2Seq) model, to improve prediction accuracy in retrosynthesis. However, these methods just utilize the sequential representations of the molecule, while ignoring the natural topological connections between atoms within the molecule. These atomic connections can provide more flexible and accurate chemical information, which is critical in many related chemical tasks like molecular representation [Duvenaud *et al.*, 2015; Gilmer *et al.*, 2017] and chemical reaction prediction [Jin *et al.*, 2017; Do *et al.*, 2019]. We believe that the absence of this molecular graph information hinders the further improvement of the present methods for retrosynthesis. How to effectively make use of this natural graphical information of the molecular structure, therefore, becomes a vital problem.

To tackle this problem, we propose Graph Enhanced Transformer(GET) framework that can enjoy the advantage of both graph-level representations and sequence-level representations. Specifically, to solve the retrosynthesis problem, we design an improved Graph Neural Network(GNN) called Graph Attention with Edge and Skip-connection (GAES) to learn each atom’s representation, and present four strategies to incorporate it with the original SMILES representation in the encoder. By fusing the natural graphical information, we can make more chemically plausible constraints on the representation learning process of molecules. Extensive experiments show that we significantly improve the prediction accuracy and the chemical-valid rate of predicted molecules. The main contributions of this paper are as follows:

- We propose a new framework called GET that fuses graphical representations with sequential representations of the target molecule to solve retrosynthesis prediction task.
- We design a powerful GNN called GAES that learns high-quality representations of atom nodes in a self-attention manner with bond features, and it is less affected by the side-effect of stacking more layers in GNN.
- We evaluate GET on USPTO-50K, a common benchmark dataset for retrosynthesis. Experimental results show that our model achieves new records for top-1 prediction accuracy among current state-of-the-art methods and especially outperforms template-free (Seq2Seq-based) methods in all tested top-*n* accuracy, demonstrating the effectiveness of fusing the molecular graph information with the SMILES sequence information.

## 2 Related Work

Prior work on retrosynthesis can be mainly summarized into two categories: template-based methods and template-free methods.

### Template-based Methods

The majority of computer-aided retrosynthetic methods in the early period were relied on encoding reaction templates or generalized subgraph matching rules. LHASA [Corey and Wipke, 1969] was the first software for retrosynthetic analysis. Recently, one of the most well-known retrosynthesis analysis tool is Synthia [Szymkuć *et al.*, 2016] that integrated about 70,000 hand-encoded reaction rules collected by manual. Based on the 60,000 reaction templates derived from 12 million single-step reaction examples, [Schreck *et al.*, 2019] introduced Reinforcement Learning (RL) into this area by treating retrosynthesis as a game whose goal is to identify policies that make (near) optimal reaction choices during each step of retrosynthetic planning. [Segler *et al.*, 2018] extracted two sets of transformation rules and combined Monte Carlo tree search with symbolic AI to discover possible retrosynthetic routes. Besides manual extracted rules, some works [Law *et* al., 2009; Segler and Waller, 2017; Coley *et al.*, 2017] tried to collect reaction templates automatically and perform retrosynthesis based on these automated templates. Although template-based methods work well in many cases, they still face a serious drawback that they generally cannot achieve accurate prediction accuracy outside of their known templates.

### Template-free Methods

Our model just belongs to this category. Emerging template-free methods are to treat retrosynthesis as a machine translation task as introduced in section 1. Since these methods do not need any reaction templates and prior chemistry knowledge, they are attracting more and more attention from academia. Moreover, without the constraint of fixed templates, they have the potential of discovering novel synthetic routes. The most related work to ours are [Zheng *et al.*, 2019] [Lin *et al.*, 2019; Karpov *et al.*, 2019; Lee *et al.*, 2019] that apply Transformer to retrosynthesis prediction.

## 3 Background

In this section, we first introduce how GNN is used in learning the molecule (or atoms) representation, and then describe how the Transformer model is previously applied in retrosynthesis prediction.

### 3.1 GNN for molecular representation learning

GNN has been widely used in learning the representation of the molecule and its atoms. Naturally, molecules can be represented as graph structure with atoms as nodes and bonds as edges. Suppose that a molecular graph *G* has initial node representations ***h***_*v*_ and edge representations ***e***_*vw*_, a typical one-layer GNN can learn new and more powerful node representations from *G* by the following message passing process described in [Gilmer *et al.*, 2017]:

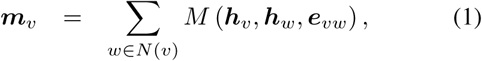

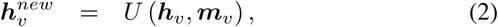

where *N*(*v*) denotes the neighbors of node *v* in graph *G, M* is the message function that is responsible for collecting information from neighbors, and *U* is the update function for fusing collected information ***m***_*v*_ with old node representation ***h***_*v*_ to obtain the new node representation 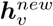. Further, we can stack several these GNN layers to capture higher-order neighbors’ information.

Then a readout function *R* can be used to integrate all node representations into a whole graph representation ***g***:

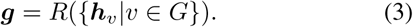

The ***h***_*v*_ and ***g***, which represent the atoms and the whole molecule, are often trained in an end-to-end way for a specific chemical task, such as chemical properties prediction, reaction prediction and molecule optimization.

### 3.2 Transformer for retrosynthesis prediction

Transformer [Vaswani *et al.*, 2017] is a Seq2Seq model that has shown excellent performance in machine translation task. Also, it has been applied in chemical reaction prediction and retrosynthesis prediction before. Given an input SMILES that represents the target molecule and a specified reaction type (optional), retrosynthesis prediction is to predict the output SMILES which represents the possible reactants that can synthesis the target molecule in the specified reaction type. Thus, retrosynthesis prediction can be deemed as a machine translation task whose source language is target molecule SMILES and the target language is reactants SMILES.

In this view, Transformer can be applied to retrosynthesis prediction as the same as to machine translation. Since Transformer is a mature model that has been widely used in natural language processing (NLP), we just give a simple introduction here. Specifically, Transformer follows an encoder-decoder structure and is composed of several combinations of multi-head attention layers and position-wise feed forward layers. The encoder consists of a stack of *N* = 6 identical layers. Each layer includes two main components: (multi-head) self-attention layer and feed-forward network. Given an input vectors (***p***_1_, …, ***p***_*n*_), ***p*** ∈ ℝ^*d*^, the *t*-th output ***s***_*t*_ of the self-attention layer is calculated by:

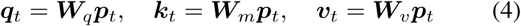

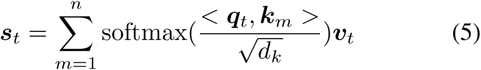

where *d*_*k*_ is the dimension of ***q*** and ***k, W***_*q*_, ***W***_*m*_, ***W***_*v*_ are weight matrices. One such operation is called one head, and we can concatenate several heads to change to multi-head self attention.

The feed-forward network is composed of two linear transformations with a ReLU activation:

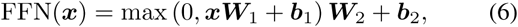

where ***W***_1_, ***W***_2_ are weight matrices and ***b***_1_, ***b***_2_ are biases.

Similarly, the decoder is also mainly composed of multi-attention layers and position-wise feed forward layers. It will generate the output SMILES step by step. At step *t*, it utilizes the encoder’s output 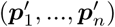 and all previous steps’ output 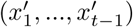 to generate the next SMILES character 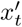. This process repeated until generating a specific termination character, i.e., 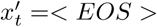.

## 4 Graph Enhanced Transformer (GET)

In this section, we provide the details about our Graph Enhanced Transformer (GET) framework for retrosynthesis prediction. Figure 1 shows an overview of GET. On the whole, GET is accord with typical encoder-decoder structure, of which the integral encoder is composed of the graph encoder and transformer encoder for learning the representation in graph-level and sequence-level respectively. Given a target molecule’s SMILES, it will first pass through the two encoders somehow to get the hidden representation of each character, and then the decoder will utilize these hidden representations to generate the output SMILES.

**Figure 1:**
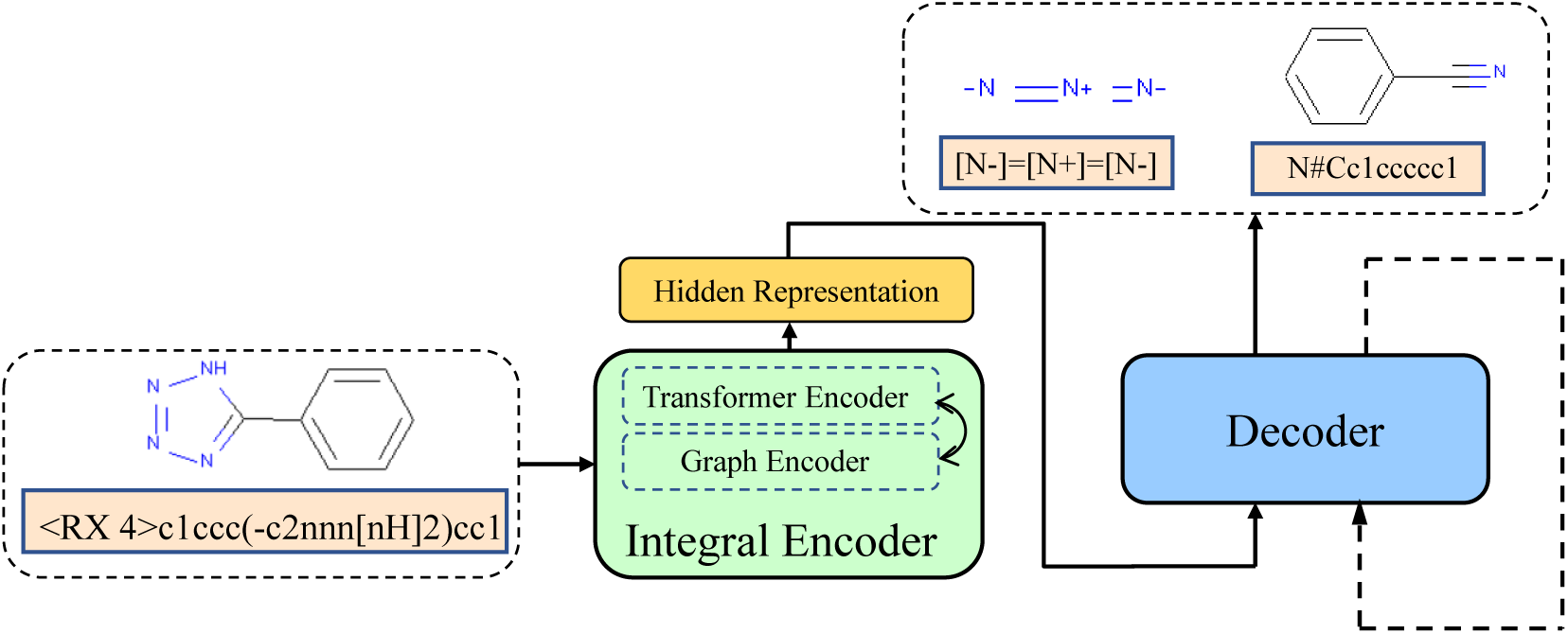
Overview of GET. For a target molecule, The input SMILES will be processed by the two-sub encoders (graph encoder and transformer encoder) somehow to be transformed to its hidden representation. Then, at each step, the decoder will utilize the hidden representation and all outputs of the previous steps to generate the present step’s SMILES character.

### 4.1 Graph Encoder

We design a new powerful GNN called Graph Attention with Edge and Skip-connection (GAES) as the graph encoder, which can learn the representation of each atom in a molecule. This graphical representation reflects the connection of atoms within a molecule and may play a significant role in further alleviating the long-term dependency problem to avoid generating chemically invalid output. We use RDKit[Landrum and others, 2006] to transform a SMILES into the molecular graph, whose nodes are atoms and edges are chemical bonds. The input representation of the atom node is a 29-dimensional vector that contains some chemical information (See Table 1) about the atom. The input representation of the edge is a 4-dimensional one-hot vector that encodes the bond types including single, double, triple and aromatic.

**Table 1:** Input representation of atom nodes.

| Atom Feature | Description |
| --- | --- |
| Atom type | C, N, O, S, P, B, F, I, K, Sn, Cl, Br, Se, Si, Zn, Cu, Mg, Pd, Pt, Li, Fe (one-hot) |
| Atom number | Numbers of protons (integer) |
| Acceptor | Accepts electrons (binary) |
| Donor | Donates electrons (binary) |
| Aromatic | In an aromatic system (binary) |
| Hybridization | sp, sp<sup>2</sup>, sp<sup>3</sup> (one-hot or null) |
| Number of Hydrogens | (integer) |

Since the SMILES sequence (*x*_1_, …, *x*_*n*_) has been transformed into a graph *G* with the input representations (***h***_1_, …, ***h***_*N*_) for nodes and {***e***_*ij*_} for edges that exist between node *i* and node *j*, our GNN will produce new representation 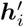 for each node *i* by the following message passing operations:

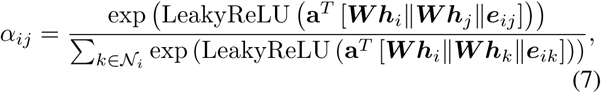

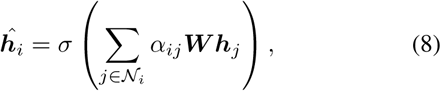

where **a** ∈ ℝ^(2*F*^′ ^+*E*)^ is a weight vector for attention mechanism and ***W*** ∈ ℝ^*F*^ *′* ^*×F*^ is a weight matrix for transforming the node features, so *F* is the input dimension of nodes, *F* ′ is the output dimension of nodes and *E* is the dimension of edges. 𝒩_*i*_ is the set of first-order neighbors of node *i* (including itself). *σ* is an activation function, e.g., ReLU.

In practice, we perform *K* multi-head attention [Veličković *et al.*, 2017] to enrich the model capacity and to stabilize the learning process. Each attention head has its own parameters and we average their outputs to get better representation:

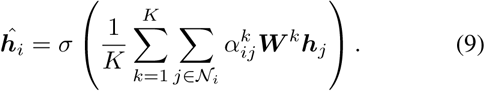

The above operations can be seen as GAT [Veličković *et* al., 2017] extended to include edge features. Then, to mitigate the accuracy reduction issue [Kipf and Welling, 2016] caused by stacked graph convolution layers, we adopt the gated skip-connection mechanism [Ryu *et al.*, 2018] to get the final representation:

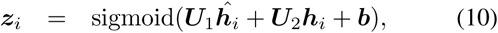

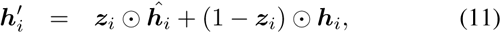

where ***U***_1_, ***U***_2_ and ***b*** are trainable parameters.

Note that the above operations are just in one layer of our GAES, and we can stack several layers to capture the information about higher-order atom neighbors so that to obtain more comprehensive representations.

### 4.2 Transformer Encoder

The transformer encoder is the same as described in section 3.2, which can capture the sequential representations of molecules (or atoms) represented by SMILES. The original SMILES (*x*_1_, …, *x*_*n*_) is changed to a sequence of vectors (***p***_1_, …, ***p***_*n*_), ***p*** ∈ ℝ^*d*^ after passing the embedding layer. And it will be further updated to vectors 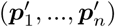 by the transformer encoder.

### 4.3 Representation Fusion

Intuitively, graphical representations reflect the intrinsic structural features of molecules and should be beneficial to generate chemical-valid and more accurate SMILES output. To make use of the graphical information, we propose four fusion strategies to fuse these graphical and sequential embeddings in the integral encoder of GET.

#### Graph Link Transformer

As shown in Figure 2 (GET-LT1), we concatenate the atom representations with embeddings of SMILES and perform a linear transformation by weight matrix ***M***. Then the new representations are sent to the transformer encoder to produce the output of the integral encoder. For non-atomic characters in SMILES, the corresponding atom representations are set to zero vector. Formally,

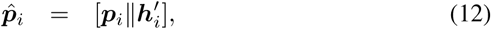

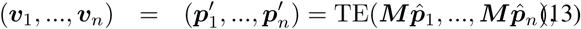

where 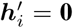 if *x*_*i*_ is non-atomic character, ‖ is the concatenation operation, (***v***_1_, …, ***v***_*n*_) is integral encoder’s output, TE is the vanilla Transformer Encoder as descirbed in 4.2.

**Figure 2:**
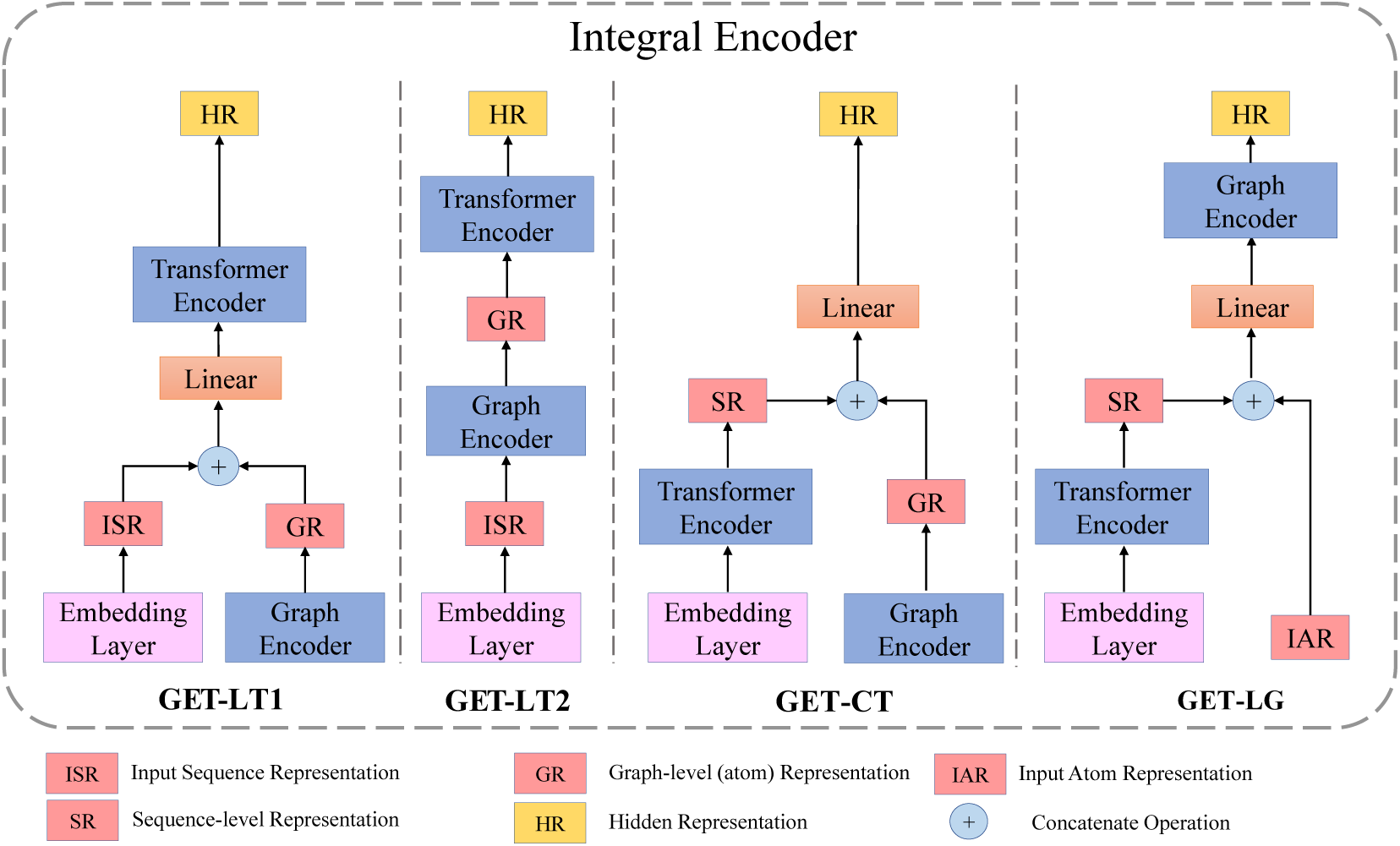
Illustration of four fusion strategies in the integral encoder of GET. The integral encoder is composed of two sub-encoders: graph encoder and transformer encoder. The embedding layer is as described in 4.2. The hidden representation of the target molecule can be obtained in four ways (GET-LT1, GET-LT2, GET-CT and GET-LG).

Considering that there may exist inconsistency between encoder and decoder since the initial atom features inputted to the graph encoder cannot be utilized by the decoder when making inference, we try another way by replacing the natural atom features (***h***_1_, …, ***h***_*n*_) with (***p***_1_, …, ***p***_*n*_) as the input representations of the graph encoder:

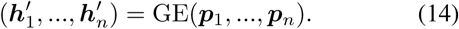

where GE is the Graph Encoder as descirbed in 4.1

In this way, the graph encoder can enhance the original sequential representations (***p***_1_, …, ***p***_*n*_) with molecule structure information directly, but the natural atom features have to be “sacrificed’. Then we send the output of the graph encoder to the transformer encoder to get the final output:

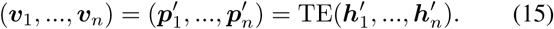

We name the first scheme GET-LT1 and the second scheme GET-LT2.

#### Graph Concatenate with Transformer

As shown in Figure 2 (GET-CT), we concatenate the outputs of the graph encoder and transform encoder, and also perform linear transformation to get the output of the integral encoder:

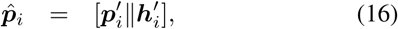

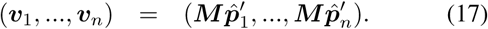

The notations are consistent with 4.3.

#### Transformer Link Graph

As shown in Figure 2 (GET-LG), the SMILES sequence first pass through the transformer encoder, then it is concatenated with natural atom features (***h***_1_, …, ***h***_*n*_) to be the input representation of the graph encoder. Those non-atomic characters are added into the molecular graph as isolated nodes which do not connect with any other node, and their “atom feature vectors” are just zero vectors. Finally, the graph encoder’s output will be the integral encoder’s output:

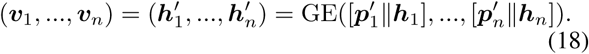

### 4.4 Decoder

The decoder is the same as vanilla Transformer’s [Vaswani *et al.*, 2017] decoder which has been introduced in section 3.2. At step *t*, the encoder’s output (***v***_1_, …, ***v***_*n*_) and all previous steps’ output 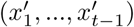 are used by the decoder to generate the next SMILES character 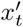 until 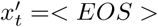.

## 5 Experiments

In this section, we evaluate our model for retrosynthesis prediction on a common benchmark dataset USPTO-50K which is derived from USPTO granted patents that includes 50,033 reactions classified into 10 reaction types. A reaction is described as a pair of sequences which consist of SMILES notations for target molecule (with reaction type) and reactants. For example, an heterocycle formation reaction is described as: (“<RX_4>c1ccc(-c2nnn[nH]2)cc1”, “N#Cc1ccccc1.[N-]=[N+]=[N-]”), where “<RX_4>” represents heterocycle formation reaction, “c1ccc(-c2nnn[nH]2)cc1” is SMILES of the target molecule, “N#Cc1ccccc1” and “[N-]=[N+]=[N-]” are SMILES of two reactants separated by “.” (as shown in Figure 1).

### 5.1 Settings

#### Dataset Splitting

Previous works [Liu *et al.*, 2017; Coley *et al.*, 2017; Zheng *et al.*, 2019; Karpov *et al.*, 2019] follow a specific split strategy with 40,029, 5,004 and 5,004 reactions for training, validation and testing, and we keep the same.

#### Implementation Details

For the graph encoder, we implement the GAES based on DGL [Wang *et al.*, 2018]. The dimension of the input node(atom) representation **h** is set to 29, and the dimension of the final output node representation **h**′ is set to 256. The number of multi-head in GAES is set to 2. To capture higher-order neighbors informaiton, we stack 3 identical GAES layers; For the transformer encoder and the decoder, we implement them using OpenNMT [Klein *et al.*, 2017]. The number of layers for the transformer encoder and decoder is set to 4, and their dimensions are set to 256. (So the final dimension of the integral encoder’s output ***v*** is also set to 256). The number of multi-head is set to 8. We use Adam with 2.0 initial learning rate to train the model and adopt 0.1 dropout rate to prevent overfitting. More detailed training settings can be found in our public code at https://github.com/papercodekl/MolecularGET.

### 5.2 Result

We compare our model with the vanilla Transformer [Vaswani *et al.*, 2017], Rule-based Expert System mentioned in [Liu *et al.*, 2017], Similarity [Coley *et al.*, 2017] and LSTM+Attention [Liu *et al.*, 2017]. Note that the results of vanilla Transformer are based on our own experiments since the results reported by previous works [Zheng *et al.*, 2019; Lin *et al.*, 2019; Karpov *et al.*, 2019; Lee *et al.*, 2019] are different from each other. Actually, their results are closed to the results of our implementation. The retrosynthesis prediction accuracy across all classes is provided in Table 2. We also test the performance of GET-LT1 when removing the reaction type from the original dataset, and the result is shown in Table 4. Besides, to further improve the accuracy, we perform the ensemble method on 10 GET-LT1 models and we provide the details and results in 5.3.

**Table 2:** Comparison of top-*n* accuracies across all classes.

| Model | top-$n$ accuracy (%) | | | |
| --- | --- | --- | --- | --- |
|  | 1 | 3 | 5 | 10 |
| Rule-based Expert System | 35.4 | 52.3 | 59.1 | 65.1 |
| LSTM+Attention | 37.4 | 52.4 | 57.0 | 61.7 |
| Similarity | 52.9 | **73.8** | **81.2** | **88.1** |
| Transformer (baseline) | 54.3 | 68.4 | 72.0 | 74.4 |
| GET-CT (ours) | 55.9 | 70.1 | 73.2 | 76.3 |
| GET-LG (ours) | 54.9 | 69.7 | 72.2 | 74.6 |
| GET-LT2 (ours) | 56.2 | 69.4 | 72.5 | 74.7 |
| GET-LT1 (ours) | **57.4** | 71.3 | 74.8 | 77.4 |

**Table 3:**
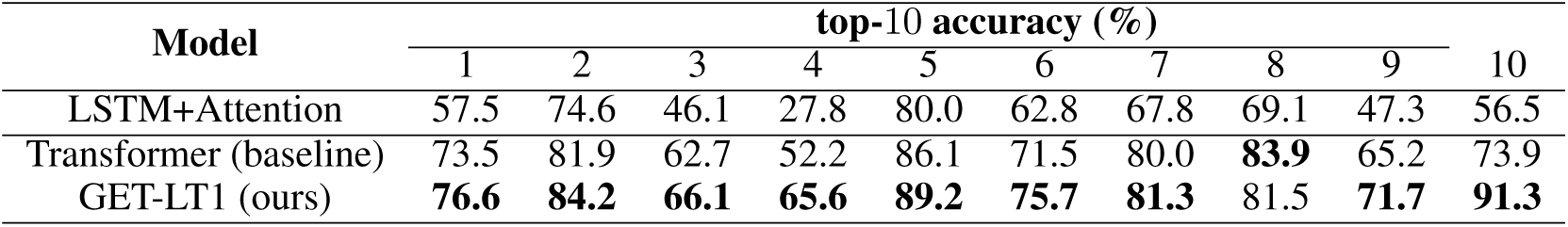
Comparison of the top-10 accuracy for each reaction class.

**Table 4:** Comparison of top-*n* accuracies across all classes without reaction type.

| Model | top-$n$ accuracy (%) | | | |
| --- | --- | --- | --- | --- |
|  | 1 | 3 | 5 | 10 |
| Similarity | 37.3 | 54.7 | **63.3** | **74.1** |
| Transformer (baseline) | 42.3 | 57.5 | 61.0 | 65.7 |
| GET-LT1 (ours) | **44.9** | **58.8** | 62.4 | 65.9 |

Results show that our models outperform all of previous methods in top-1 accuracy, and our best model GET-LT1 achieves the new state-of-the-art among all template-free (Seq2Seq-based) methods, i.e, LSTM+Attention, vanilla Transformer and models of GET. Compared with vanilla Transformer, GET-LT1 can improve the prediction accuracy by 3.1%, 2.9%, 2.8% and 3.0% in top-1, top-3, top-5 and top-10 accuracy. Other variants also have varying degrees of improvement over vanilla Transformer. The reasons why GET-LT1 performs best may be as follows: (1) Compared with GET-LT2, GET-LT1 utilizes the atom information. (2) Compared with GET-LG, the learned representations by GET-LT1 are more determined by the transformer encoder. Since the decoder of GET is a “sequential decoder”, it may be better to give the “sequential encoder”(i.e., transformer encoder) more weights while using the graphical encoder (i.e., graph encoder) as an auxiliary enhancement. Thus, letting the representations first pass through the graph encoder and then pass through the Transformer encoder can obtain better results. (3) Compared with GET-CT, GET-LT1 fuses the graphical and sequential representations in a cascaded way while not in a parallel way (i.e., concatenation). Since the concatenation operation cannot distinguish the importance of two representations as cascaded way does (the latter is more important as explained in (2)), it performs worse.

Moreover, as shown in Table 4, our model can retain this comprehensive superiority after removing the reaction type, demonstrating that molecule structure information, which brings more chemically constraints, can help Transformer to predict more accurate reactants. In addition, we present the detailed top-10 accuracy of three Seq2Seq-based models (LSTM+Attention, vanilla Transformer and GET-LT1) for each reaction class in Table 3. Results show that our approach can improve vanilla Transformer on 9 of 10 reaction classes by a margin of 1.3% to 17.4%, indicating the better generalization ability and comprehensiveness of our model.

Furthermore, the rate of producing chemical-invalid SMILES for different beam sizes are shown in Table 5 (with reaction type). As can be seen, after fusing molecule structure information, the model is more inclined to generate chemical-valid SMILES compared with vanilla Transformer, since the graphical representations, which directly capture the topological connection of atoms, are able to break the limitation of the SMILES sequence and can give the model additional guidance to produce chemical-valid compounds.

**Table 5:** The rate of producing chemical-invalid SMILES for different beam sizes.

| Model | invalid SMILES' rate (%)<br>when beam size $k =$ | | | |
| --- | --- | --- | --- | --- |
|  | 1 | 3 | 5 | 10 |
| LSTM+Attention | 12.2 | 15.3 | **18.4** | **22.0** |
| Transformer (baseline) | 3.5 | 14.3 | 20.3 | 30.2 |
| GET-LT1 (ours) | **2.2** | **13.4** | 19.5 | 29.3 |

### 5.3 Ensemble for better performance

To obtain more accurate results, we overall consider 10 models’ results by the ensemble method. Specifically, in the vanilla Transformer or GET, the decoder output the predicted character *x*_*t*_ based on the probability vector **z**_**t**_. The *i*-*th* dimension of **z**_**t**_ represents the probability of outputing the *i*-*th* character in the vocabulary. Suppose that 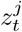 is the *j*-*th* model’s probability vector at step *t*, we average all of the 10 models’ probability vectors to get the averaged probability vector 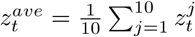. Instead, we use 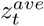 to select *x*_*t*_.

For GET-LT1, we choose 10 models which are from 410,000 to 500,000 training steps (10,000 as the interval) to ensemble. For Transformer, we choose 10 models from 110,000 to 200,000 training steps. We experiment 3 times with different random seeds (41, 42, 43) and the average results are shown in Table 6.

**Table 6:** Comparison of top-*n* accuracies with the reaction type under the ensemble method (average results).

| Model | top-$n$ accuracy (%) | | | |
| --- | --- | --- | --- | --- |
|  | 1 | 3 | 5 | 10 |
| Transformer (baseline) | 56.1 | 71.1 | 75.2 | 78.2 |
| GET-LT1 (ours) | **59.1** | **73.4** | **76.4** | **78.7** |

## 6 Conclusion and Future Work

We propose Graph Enhanced Transformer(GET), an effective framework that successfully combines the graphical and sequential representations of the molecule to improve the retrosynthesis prediction performance. Experiments indicate that our model outperforms state-of-the-art template-free methods on USPTO-50K dataset, and shows promising ability in reducing chemical-invalid SMILES rate.

## References

Connor W Coley, Luke Rogers, William H Green, and Klavs F Jensen. Computer-assisted retrosynthesis based on molecular similarity. ACS central science, 3(12):1237–1245, 2017.

EJ Corey and W Todd Wipke. Computer-assisted design of complex organic syntheses. Science, 166(3902):178–192, 1969.

Kien Do, Truyen Tran, and Svetha Venkatesh. Graph transformation policy network for chemical reaction prediction. In Proceedings of the 25th ACM SIGKDD International Conference on Knowledge Discovery & Data Mining, pages 750–760. ACM, 2019.

David K Duvenaud, Dougal Maclaurin, Jorge Iparraguirre, Rafael Bombarell, Timothy Hirzel, Alán Aspuru-Guzik, and Ryan P Adams. Convolutional networks on graphs for learning molecular fingerprints. In Advances in neural information processing systems, pages 2224–2232, 2015.

Justin Gilmer, Samuel S Schoenholz, Patrick F Riley, Oriol Vinyals, and George E Dahl. Neural message passing for quantum chemistry. In Proceedings of the 34th International Conference on Machine Learning-Volume 70, pages 1263–1272. JMLR. org, 2017.

Wengong Jin, Connor Coley, Regina Barzilay, and Tommi Jaakkola. Predicting organic reaction outcomes with weisfeiler-lehman network. In Advances in Neural Information Processing Systems, pages 2607–2616, 2017.

Pavel Karpov, Guillaume Godin, and Igor V Tetko. A transformer model for retrosynthesis. In International Conference on Artificial Neural Networks, pages 817–830. Springer, 2019.

Thomas N Kipf and Max Welling. Semi-supervised classification with graph convolutional networks. arXiv preprint 1609.02907, 2016.

Guillaume Klein, Yoon Kim, Yuntian Deng, Jean Senellart, and Alexander M Rush. Opennmt: Open-source toolkit for neural machine translation. arXiv preprint 1701.02810, 2017.

Greg Landrum et al. Rdkit: Open-source cheminformatics, 2006.

James Law, Zsolt Zsoldos, Aniko Simon, Darryl Reid, Yang Liu, Sing Yoong Khew, A Peter Johnson, Sarah Major, Robert A Wade, and Howard Y Ando. Route designer: a retrosynthetic analysis tool utilizing automated retrosynthetic rule generation. Journal of chemical information and modeling, 49(3):593–602, 2009.

AA Lee, Q Yang, V Sresht, P Bolgar, X Hou, JL Klug-McLeod, and CR Butler. Molecular transformer unifies reaction prediction and retrosynthesis across pharma chemical space. Chemical communications (Cambridge, England), 2019.

Kangjie Lin, Youjun Xu, Jianfeng Pei, and Luhua Lai. Automatic retrosynthetic pathway planning using template-free models. arXiv preprint 1906.02308, 2019.

Bowen Liu, Bharath Ramsundar, Prasad Kawthekar, Jade Shi, Joseph Gomes, Quang Luu Nguyen, Stephen Ho, Jack Sloane, Paul Wender, and Vijay Pande. Retrosynthetic reaction prediction using neural sequence-to-sequence models. ACS central science, 3(10):1103–1113, 2017.

Seongok Ryu, Jaechang Lim, Seung Hwan Hong, and Woo Youn Kim. Deeply learning molecular structure-property relationships using attention- and gate-augmented graph convolutional network. arXiv preprint 1805.10988, 2018.

John S Schreck, Connor W Coley, and Kyle JM Bishop. Learning retrosynthetic planning through self-play. arXiv preprint 1901.06569, 2019.

Marwin HS Segler and Mark P Waller. Neural-symbolic machine learning for retrosynthesis and reaction prediction. Chemistry–A European Journal, 23(25):5966–5971, 2017.

Marwin HS Segler, Mike Preuss, and Mark P Waller. Planning chemical syntheses with deep neural networks and symbolic ai. Nature, 555(7698):604, 2018.

Sara Szymkuć, Ewa P Gajewska, Tomasz Klucznik, Karol Molga, Piotr Dittwald, Michal Startek, Michal Bajczyk, and Bartosz A Grzybowski. Computer-assisted synthetic planning: The end of the beginning. Angewandte Chemie International Edition, 55(20):5904–5937, 2016.

Ashish Vaswani, Noam Shazeer, Niki Parmar, Jakob Uszkoreit, Llion Jones, Aidan N Gomez, Łukasz Kaiser, and Illia Polosukhin. Attention is all you need. In Advances in neural information processing systems, pages 5998–6008, 2017.

Petar Veličković, Guillem Cucurull, Arantxa Casanova, Adriana Romero, Pietro Lio, and Yoshua Bengio. Graph attention networks. arXiv preprint 1710.10903, 2017.

Minjie Wang, Lingfan Yu, Quan Gan, D. Zheng, Yu Gai, Zihao Ye, Mufei Li, Jinjing Zhou, Qi Huang, Junbo Zhao, Haibin Lin, Chao Ma, Damon Deng, Qipeng Guo, Hao Zhang, Jinyang Li, Alexander J Smola, and Zheng Zhang. Deep graph library, 2018.

David Weininger. Smiles, a chemical language and information system. 1. introduction to methodology and encoding rules. Journal of chemical information and computer sciences, 28(1):31–36, 1988.

Shuangjia Zheng, Jiahua Rao, Zhongyue Zhang, Jun Xu, and Yuedong Yang. Predicting retrosynthetic reaction using self-corrected transformer neural networks. arXiv preprint 1907.01356, 2019.

